# Cis-Regulatory Accessibility Directs Muller Glial Development and Regenerative Capacity

**DOI:** 10.1101/752428

**Authors:** Leah S. VandenBosch, Stefanie G. Wohl, Matthew S. Wilken, Kristen Cox, Laura Chipman, Thomas A. Reh

**Affiliations:** Department of Biological Structure, University of Washington, Box 357420, Seattle, WA, 98195, USA; Molecular and Cellular Biology Program, University of Washington, Seattle, WA, USA; Institute for Stem Cells and Regenerative Medicine, University of Washington, Box 358056, Seattle, WA, 98109, USA; Department of Biological and Vision Sciences, The State University of New York, College of Optometry, New York, NY, USA

## Abstract

Diseases and damage to the retina lead to losses in retinal neurons and eventual visual impairment. Although the mammalian retina has no inherent regenerative capabilities, fish have robust regeneration from Müller glia (MG). Recently, we have shown that driving expression of Ascl1 in adult mouse MG stimulates neurogenesis similar to fish regeneration. The regeneration observed in the mouse is limited in the variety of neurons that can be derived from MG; Ascl1-expressing MG primarily generate bipolar cells. To better understand the limits of MG-based regeneration in mouse retinas, we used ATAC- and RNA-seq to compare newborn progenitors with MG. Our analysis demonstrated striking similarities between MG and progenitors, with losses in regulatory motifs for neurogenesis genes. Young MG were found to have intermediate expression profiles and accessible DNA, which is mirrored in the ability of Ascl1 to direct bipolar neurogenesis in young MG. When comparing what makes bipolar and photoreceptor cells distinct from glial cells, we find that bipolar-specific accessible regions are more frequently linked to bHLH motifs and Ascl1 binding, indicating that Ascl1 preferentially binds to bipolar regions. Overall, our analysis indicates a loss of neurogenic gene expression and motif accessibility during glial maturation that may prevent efficient reprogramming.

## Introduction

Neuron loss caused by disease and damage to the mammalian retina can lead to permanent vision loss. While some species are readily capable of regenerating lost neurons, mammalian retinas are not regenerative. In the mammalian retina, neuron loss caused by direct damage to the retina leads to reactive gliosis of the Müller glia (MG), similar to that of astrocytes in the brain (Bringmann et al. 2009).

Teleost fish, by contrast, are capable of regenerating retinal neurons, including photoreceptors and ganglion cells, after damage. This regeneration is carried out by the MG, which respond to damage by generating progenitor-like cells, similar to those in the developing retina (Goldman 2014; Gemberling et al. 2013). This regeneration is accompanied by waves of gene expression and morphological changes to the MG, regulated by epigenomic changes directing regeneration (for review see VandenBosch and Reh 2019). The murine retina also undergoes epigenomic changes after damage, but neurogenic programs are not re-expressed, and neuronal regeneration does not occur (VandenBosch and Reh 2019).

A critical difference between the fish MG and the mammalian MG in their response to damage is in their expression of the proneural transcription factor Ascl1. In fish, Ascl1 is quickly upregulated after damage, and is necessary for regeneration of new neurons (Ramachandran et al. 2010; Fausett et al. 2008). In the murine retina, Ascl1 is expressed in retinal progenitors and necessary for development of rods and bipolar cells (Ohsawa and Kageyama 2008); however it is not expressed in mature MG and after damage or in disease models, mouse MG do not spontaneously upregulate Ascl1 (Bringmann et al. 2009; Ueki et al. 2015). We recently directed Ascl1 expression to mouse MG with a tetO transgenic approach to test whether Ascl1 expression is sufficient to induce regeneration. Expression of Ascl1 in the MG of mice over the age of 12 days post-natal (P) stimulated MG to generate new bipolar neurons after NMDA damage (Ueki et al. 2015). In adult mice, however, Ascl1 over-expression in the MG is no longer sufficient to induce neurogenic potential, even in the presence of damage (Jorstad et al. 2017). In mature mice, the addition of the histone deacetylase trichostatin-A (TSA), in combination with Ascl1 overexpression and NMDA damage is required for neurogenesis; up to 30% of the Ascl1-expressing MG produce functional bipolar- and amacrine-like interneurons, confirmed by single cell transcriptomics, electrophysiology, and electron microscopy (Jorstad et al. 2017).

The fact that the neurogenic program can be activated in mature MG by Ascl1 only in combination with HDAC inhibition suggests that epigenetic mechanisms may limit regeneration from the MG. In addition, even with the addition of HDAC inhibitors, the Ascl1-expressing MG only generate a subset of the neurons in the retina, suggesting that epigenetic factors may also limit the types of neurons that can be regenerated from mammalian MG. Thus, the expression of Ascl1, along with HDAC inhibition, does not fully recapitulate the multipotent progenitor state present in developing retina. Therefore, we asked whether changes in the transcriptome or epigenetic landscape might account for the difference in neurogenic potential between MG and retinal progenitors.

To address the question of what distinguishes mature MG from late progenitors, we performed a transcriptomic and epigenomic comparison of FACS-isolated postnatal progenitors, young MG (before eye-opening at P12), and adult MG. Analysis by ATAC and RNA sequencing demonstrates a clear trend in the loss of neurogenesis-related motif accessibility and expression. Immature MG are found to have an intermediate epigenomic and transcriptomic profile. To test whether the intermediate profile of young MG regulates their neurogenic potential, we over-expressed Ascl1 in developing MG and have identified key restriction points in the neurogenic potential of MG that correlate with changes in the accessible chromatin landscape.

## Results

### Chromatin Accessibility in Retinal Progenitors

To determine the differences in the epigenomic landscape of retinal progenitors and developing MG, we used an Assay for Transposase-Accessible Chromatin (ATAC) sequencing to probe for differences in accessibility (Figure 1A). To isolate retinal progenitor cells, we used a knock-in Sox2-GFP mouse line that expresses GFP under control of the Sox2 promoter (Arnold et al. 2011). The great majority of Sox2+ cells at this age are retinal progenitors, though there is a small population of Sox2+ amacrine cells that can be distinguished from the progenitors by their high level of GFP. The retinas of P2 pups were dissociated into single cells and the GFP+ cells were sorted by Fluorescence-Activated Cell Sorting (FACS); the small number of strongly fluorescent amacrine cells were sorted separately from the more abundant progenitors (Figure S1). Sorted cells were used for two runs of ATAC-seq, reads were mapped and peaks were called using HOMER (Heinz et al. 2010) findPeaks. Two biological replicates were carried out (first run of 121M reads and 101k peaks, with a replicate with 95M reads and 68k peaks). When overlapped by BEDOPS (Neph et al. 2012), there were 40k common peaks between the samples, which were used for the subsequent analysis.

**Figure 1.**
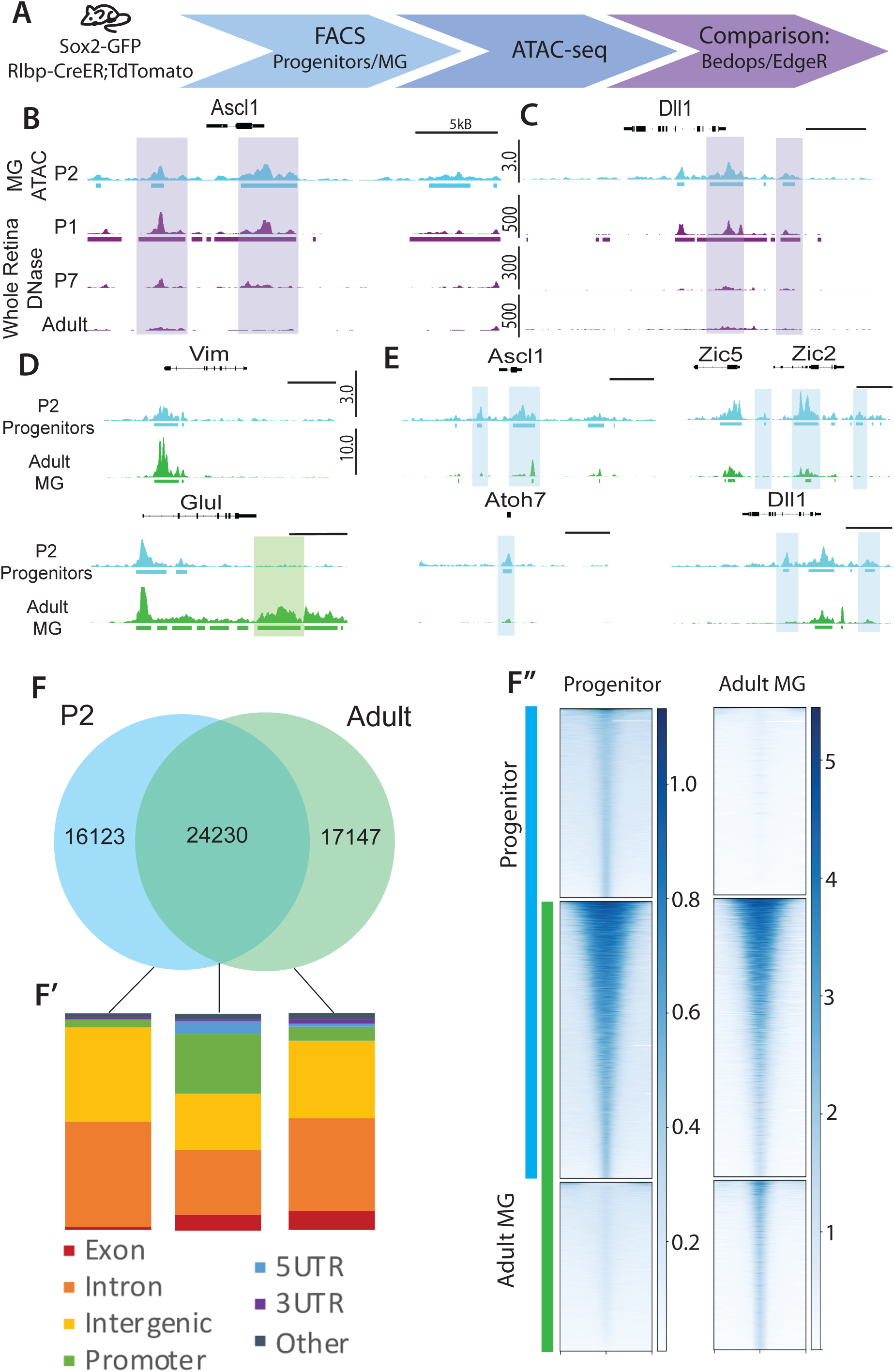
Progenitor and Müller Glial accessibility is generally similar. A. ATAC experimental design. B-C. Genomic tracks of P2 progenitor ATAC and whole retina DNase(Wilken et al. 2015) around Ascl1 and Dll1. D. Tracks for progenitors and adult MG (Jorstad et al. 2017) ATAC-seq at glial genes and E. progenitor genes. F. BEDOPS overlaps of progenitor and adult MG peak calls. F’. Genomic region annotation of accessibility profiles. F’’. Tag density in progenitor and adult MG ATAC at accessibility profiles.

We previously characterized chromatin accessibility by DNase I in whole retinas at P0, P7 and Adult (Wilken et al. 2015). We compared the progenitor ATAC results with that from DNaseI-seq of the whole retina at the three ages. At P0, approximately 30% of the retinal cells are progenitors, while at P7 there are few progenitors remaining at the retinal periphery and none in the adult (Young 1985). Thus, we would anticipate the greatest overlap in accessible peaks between the progenitor ATAC-seq and the P0 retina. We found that progenitor-specific genes, such as Ascl1, show similar accessibility in the P0 retina and P2 FACS-purified progenitor cells (Figure 1B,C). Comparison of the progenitor ATAC-seq with older P7 and adult whole retina DNaseI-seq showed a reduction in accessibility at regions near progenitor-specific genes, consistent with the loss of progenitor cells as the retina matures.

### Chromatin Landscapes in MG and Progenitors Have a High Level of Overlap

We compared the progenitor ATAC-seq data to our previously published mature adult MG ATAC-seq to determine if there are specific molecular differences in chromatin accessibility between these two cell types. The accessible chromatin in MG peaks was assessed in both FACS purified MG from adult mouse retina, as well as from MG maintained in dissociated culture. In previous studies, we found that a small amount of rod photoreceptor DNA and RNA is carried along with the adult MG during FACS (Jorstad et al. 2017); however, when we maintain the MG in dissociated culture, very few rods (<0.1%) survive. Therefore, to reduce the contribution from rod contamination in our downstream analyses, MG-specific regions of accessibility were chosen that overlapped with DNase-seq from cultured P12 MG(Ueki et al. 2015). When we examine the peaks that are present in both the freshly isolated MG ATAC-seq and the peaks from the cultured MG, we find they largely overlap (Figure S2).

When we then compared the MG accessible peaks with those of the progenitor cells, we find that there is extensive overlap in overall accessibility. Approximately sixty percent of the progenitor peaks (24,230 peaks) were shared between progenitors and MG, while P2 progenitors had16k exclusive peaks, and the adult MG have 17k unique peaks (Figure 1F). Analysis of the peaks that are unique to either progenitor cells or MG allowed some general conclusions: (1) Both progenitor cells and MG have similar regions of accessible chromatin near genes that are expressed at high levels in MG; however, the MG show a greater signal at many of these peaks, and also had unique peaks near glial genes that were not present in the progenitor cells (Figure 1D). (2) Somewhat surprisingly, genes that are important for progenitor cell function (eg. Ascl1, Dll1) had many of the same accessible regions near promoters in both the MG and the progenitors (Figure 1E), though in many cases the P2 progenitors had more regions of accessibility at these genes than the mature MG. The difference in accessibility between these cell types varies from reduced peak height to a complete loss of some peaks. (3) Promoter regions were over-represented in regions of shared accessibility, while peaks that were unique to either MG or progenitor cells were predominantly found in intronic and intergenic regions (Figure 1F;F’). Read density in these categories of accessibility showed that shared accessible regions had an overall higher read density in broader regions, whereas cell type specific accessible regions had narrow regions of read density (Figure 1F’’). These differences suggest that while promoter and major regulatory regions retain similar accessibility, many putative regulatory regions differed in accessibility between these cell types, likely reflecting the difference in their respective patterns of gene expression.

### Cis-Regulatory Binding Sites Characterize Glial Development

In order to explore putative regulatory regions that are specific to progenitor and MG populations, we analyzed and annotated regions of changing accessibility. We defined regions of changing accessibility as regions that were either unique peaks called by Homer and overlapped by BEDOPS (Neph et al. 2012) from P2 to Adult, or alternatively showed differences in tag density by a logFC (fold change) > 2 in the top 1000 regions of Differential Accessibility (DA). These two different pipelines gave similar results for gene ontology and binding motif annotation (Tables S1, S2).

When we carried out gene ontology (GO) analysis for those regions that had greater accessibility in the progenitor cells than in the MG (Loss of Accessibility or LOA), we found these regions were associated with genes that were enriched for GO terms of Neural Development/Neurogenesis and Developmental Process/Cell Differentiation (Figure 2A). By contrast, the peaks that were not accessible in retinal progenitors, but present in mature MG (ie. Gain of Accessibility or GOA), were associated with genes that were enriched for GO terms of more general cell function: e.g. metabolic and cell process genes (Figure 2B). Those accessible regions present in both cell types are also enriched primarily in metabolic genes (Table S1).

**Figure 2.**
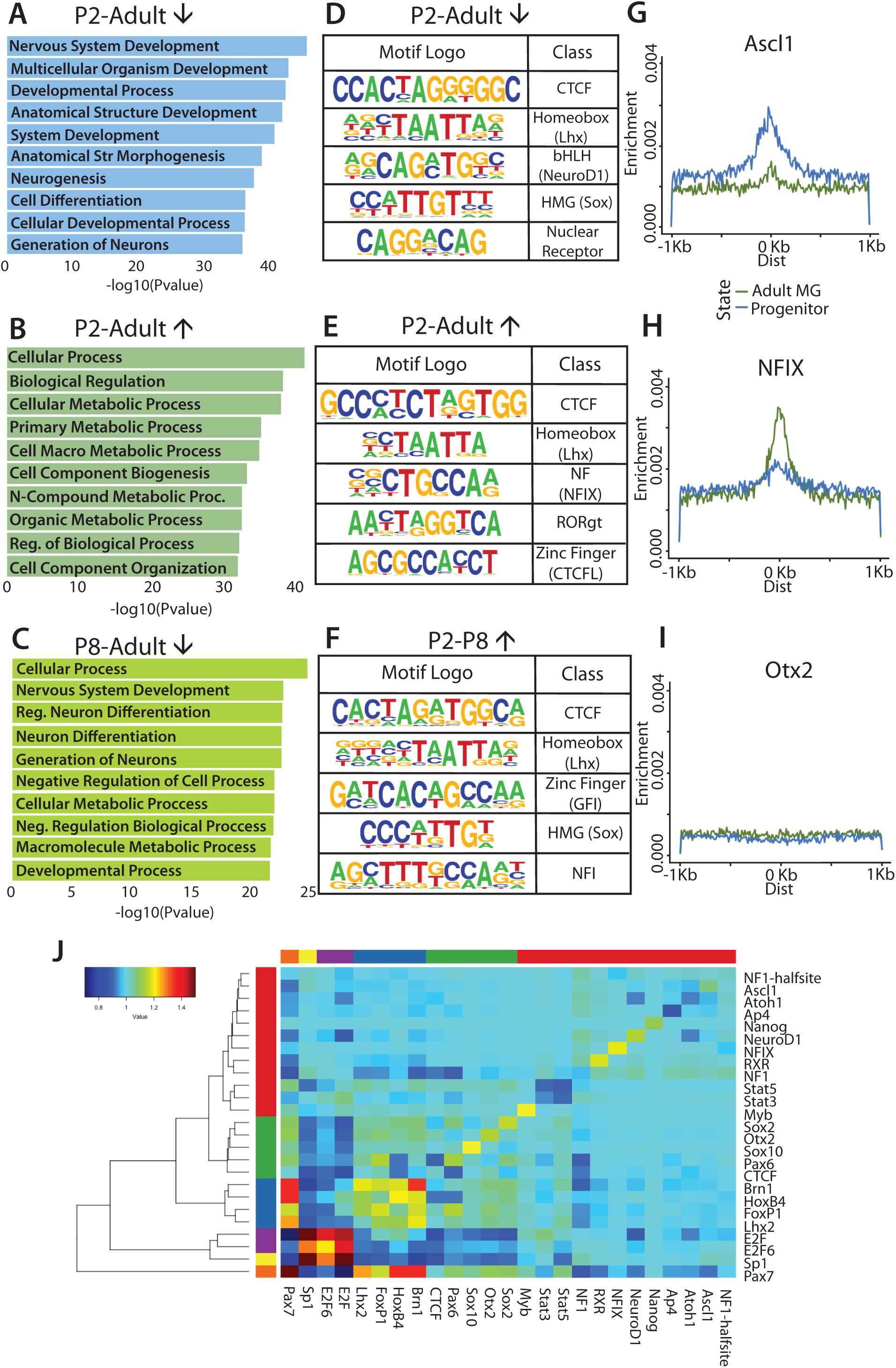
Cis-regulatory binding motifs differ in Müller glia development. A-C. Gene ontology associated with accessibility changes (A. P2-adult decreasing, B. P2-Adult increasing, C. P8-Adult decreasing,). D-F. Predicted motif enrichment for accessibility changes (D. P2-Adult decreasing [bedops], E. P2-Adult increasing [bedops], F. P2-P8 increasing [DA]). G-I. Central enrichment of motifs for progenitor- and adult MG-specific accessibility regions. J. Co-occurrence analysis for top predicted motifs in progenitor-specific accessibility regions clustered by co-occurrence ratios.

We next performed ATAC-seq on FACS-purified immature P8 MG to better understand the dynamic changes in the accessible chromatin that occur as MG mature (Figure S3). For the P8 sample, there were 49.6M reads and 16k peaks by Homer findPeaks, of which 98.35% overlapped with the P7 DNaseI whole retina dataset. We carried out DA analysis between the P8 and adult MG, and the progenitor cells and P8 MG. Interestingly, we found GO terms of Nervous System Development and Generation of Neurons as some of the top Biological Process terms in the P8 MG accessible regions that are lost or reduced in the mature MG cells (Figure 2C). Thus, it appears that many of the putative cis-regulatory regions near genes associated with neurogenesis are still accessible in the immature P8 MG, and suggests these cells may be more amenable to reprogramming than mature MG.

The peaks of accessible chromatin near genes associated with neurogenesis that are unique to progenitor cells (ie. not present or reduced in the mature MG) may be relevant to the differences between these cells in their ability to generate neurons. We analyzed these putative neurogenesis-related cis-regulatory elements for transcription factor binding motifs using Homer findMotifsGenome against randomly selected background regions. We found that the top five transcription factor motifs present in the peaks unique to progenitor cells (LOA) were - CTCF, Lhx (eg. Lhx2), bHLH (eg. NeuroD1), HMG (eg. Sox2) and nuclear receptor (Figure 2D). Progenitor cells are known to express several members of each of these transcription factor families; for example, the proneural bHLH factors Neurog2, Olig 2 and Ascl1 are all expressed in retinal progenitors at this stage (Brzezinski et al. 2011). When comparing Eboxes with specificity to Ascl1 and Neurog2, we found that Ascl1-specific Eboxes had far more distinct central enrichment, especially in regions of progenitor-specific accessibility, whereas Neurog2 Eboxes are centrally depleted (Figure S4A,B). Factors such as Lhx and Sox are known to be expressed in both progenitors and MG, and thus their presence in progenitor-specific accessible domains is likely indicative of restructuring of regulatory regions during development.

While the accessible chromatin regions specific to progenitors were enriched for proneural transcription factors, the accessible regions present in the MG, but not present in the progenitors (GOA), have a very different set of enriched transcription factor binding motifs. Although CTCF and Lhx motifs were still among the top 5, the NFI (eg. NFIx) binding motif is enriched in those accessible regions unique to the MG (Figure 2E). This motif was also present in the accessible regions that differ between the immature P8 MG and the progenitors, making this an early marker of the unique MG epigenetic landscape (Figure 2F). The motifs enriched in the MG-specific accessible chromatin regions may well reflect the importance of Lhx and NFI transcription factors in MG maturation(Hägglund et al. 2011; de Melo et al. 2016; Gordon et al. 2013; Shu et al. 2003)

Central enrichment modeling also highlighted the differences between progenitors and MG. For example, progenitor-specific accessible regions were centrally enriched for predicted Ascl1 motifs, with much lower accessibility in the adult MG at these sites (Figure 2G). The opposite is true for the NFIx motif: adult MG-specific accessibility profile showed much higher enrichment for NFIx binding motifs within accessible domains than the progenitors (Figure 2H). By contrast, homeobox domains not enriched in any of these regions (e.g. Otx2) showed no central enrichment for either progenitor- or MG-specific accessible domains (Figure 2I). These motif enrichments further support a role bHLH factors in maintaining the neurogenic potential in progenitor cells, and of NFI transcription factors in regulating MG fate.

Since many DNA-binding transcription factors may act in tandem with other transcription factors in a combinatorial manner to regulate gene expression, we analyzed co-occurrence between the top predicted binding motifs for progenitor (Figure 2J) and adult MG (Figure S4C) accessible domains. Through this analysis, we found that Sp1 and E2F domains commonly co-occur in both progenitor- and MG-specific accessible regions. We also found some co-occurrence between Brn1, HoxB4, FoxP1, and Lhx2 motifs, particularly in progenitor-specific accessible domains. Overall, co-occurrence analysis does not indicate any particular co-regulatory group that might interact with the major motifs that are associated with progenitor (Ascl1) or glial (Nfix) cell fate.

### Immature MG Expression is Intermediate to Progenitors and Mature MG

To better understand the molecular basis for the difference in accessibility between the progenitors and the MG, we carried out RNA-seq on FACS purified progenitors and immature and mature MG (Figure 3A). MG were sorted using Rlbp-creER:flox-stop-tdTomato reporter mice, and progenitors were sorted using Sox2-GFP mice as previously described. Differential expression analysis was performed to demonstrate overall differences between progenitors and mature MG. We selected the 1000 most highly expressed genes with logFC >2, which were differentially expressed in MG (Gain of Expression, GOE) or in progenitors (Loss of Expression, LOE) by logCPM (counts per million) values (Figure 3B). Genes were clustered via k-means clustering, and GO terms were associated with specific clusters (Figure 3D, Supplementary Tables 4, 5). Genes that showed marked increases over the period from P2 to adult MG had Biological Process terms of Response to Stimulus, and Biological Regulation, while genes that were highly expressed in progenitors, but expressed at much lower levels in MG were associated with GO terms that included Cell Cycle, Nervous System Development and Neurogenesis (Figure 3D, Supplementary Table 5). Thus, the changes in gene expression between progenitors and MG are very similar to the changes in DNA accessibility.

**Figure 3.**
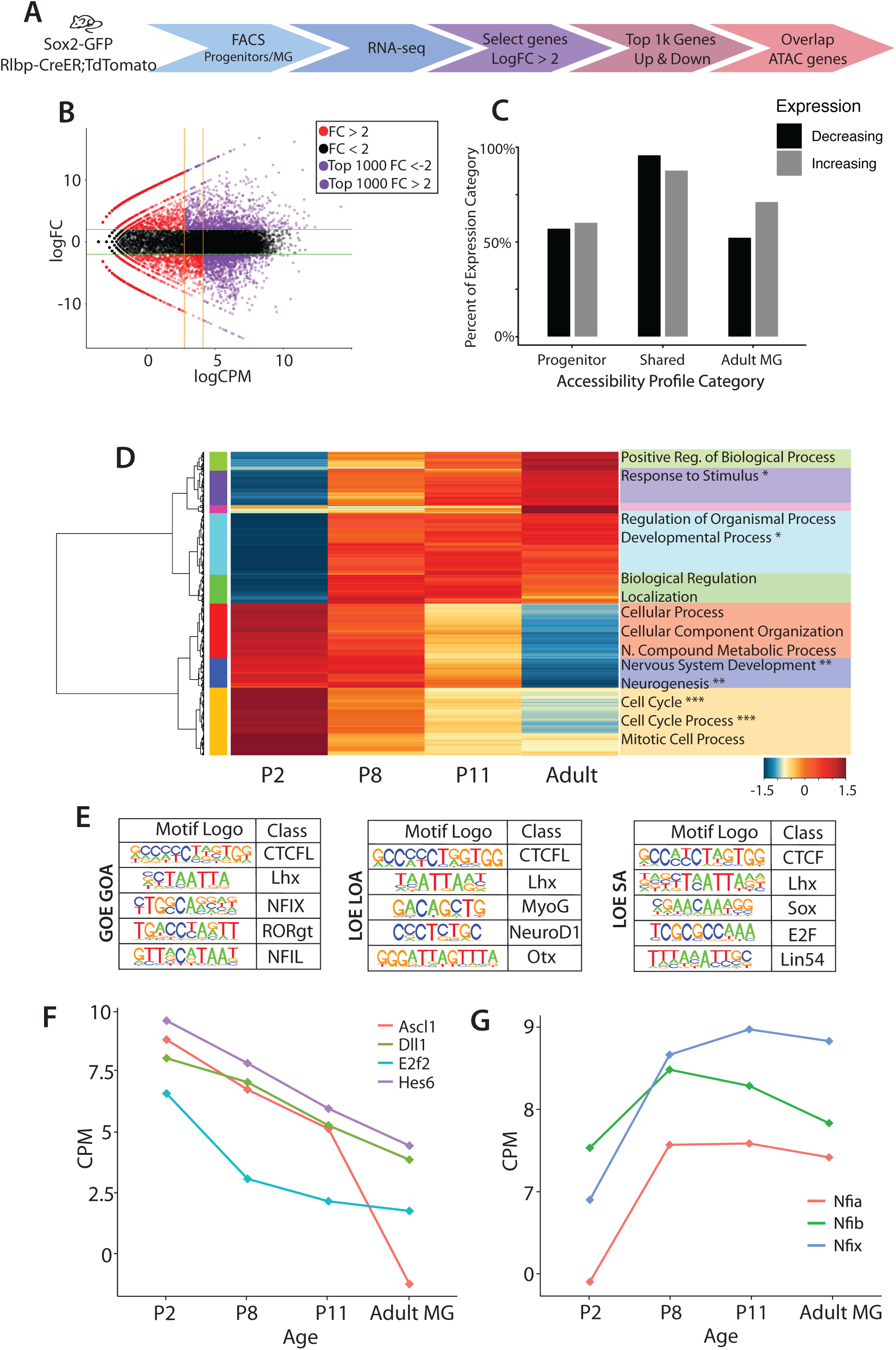
Glial expression and accessibility profiles follow developmental trends. A. Experimental design for FACS isolation and RNA-seq of developing MG. B. MA plot of expression from progenitors to adult MG demonstrating thresholds for filtering high fold change and counts for the top genes with changing expression. C. Overlaps of top gene expression changes with ATAC profiles. The percentage of genes that increase or decrease expression are represented on the y axis, split by ATAC categories(x). D. Euclidian hierarchical clustering by expression changes from P2 to Adult, heat map of top gene expression, and related gene ontology categories. Categories marked with a * indicates categories that overlap with ATAC MG increased accessibility. Those marked with a ** indicate where progenitor-specific accessibility overlaps, and *** indicates where shared accessibility overlaps. E. Motif enrichment for accessible regions with gained expression and accessibility (GOE GOA), lost expression and accessibility (LOE LOA), and lost expression and shared accessibility (LOE SA). F-G. Expression profiles of (F) progenitor genes of interest, and (G) NFI transcription factors.

We also compared the gene expression in the immature MG at P8 and P11. These immature MG expressed many of the genes that are expressed in the retinal progenitors, albeit at a lower level than the progenitors, but in addition they expressed genes more characteristic of mature MG (Figure 3D). The heatmap shows that four of the five clusters of “MG-specific” genes had the greatest increase between P2 and P8. Genes that lost expression most rapidly, in the tan cluster are primarily associated with the cell cycle. In contrast, the cluster (blue) most closely associated with the GO terms of Neurogenesis and Nervous System Development declined over the first postnatal week more gradually, but had very low levels of expression in mature MG (Figure 3D). Comparing expression up and down across MG maturation revealed an intermediate level of expression in young MG with progenitor gene expression retained at relatively high levels through P8.

Are the changes in gene expression between MG and retinal progenitors reflected in their chromatin accessibility? To answer this question, we compared the genes that change in their expression (GOE, LOE) with the developmental ATAC categories (GOA, LOA). We directly compared the GOE and LOE genes to accessibility categories (GOA, LOA, SA [Shared Accessibility]) using gene annotations by GREAT to neighboring genes, yielding a percent of total genes for GOE and LOE categories for each accessibility profile (Figure 3C, Supplementary Table 6). The best correlation between changes in gene expression and changes in accessibility was found in the adult MG: GOA regions are primarily associated with genes that increased in expression (Figure 3C, right hand bars). This correlation was not as clear in the other categories. Accessible chromatin regions that were shared between progenitors and MG were associated with genes that showed a small reduction in expression between progenitors and MG. However, accessible chromatin regions that were specific to progenitors were not necessarily associated with genes that were more highly expressed in progenitors than MG. Despite the lack of good correlation between accessibility and gene expression on a global level, analysis of specific genes and sets of genes are informative. For example, the regions of lost expression and accessibility (LOE/LOA) are best associated with Neurogenic GO categories, and this trend is continued in comparisons from P8 to Adult MG LOE/LOA (Supplementary Table 7).

By overlapping the ATAC-seq data with the gene expression results, we were able to better define some of the differences in transcription factor expression and putative binding at gene-associated regions that might be regulating the difference in neurogenic competence between these two cell types. For example, LOE/LOA regions were enriched for binding motifs for proneural bHLH factors, whereas those genes that gain accessibility and expression (GOE/GOA) were enriched for binding motifs for NFI and ROR (Figure 3E). These results are in line with those obtained from the analysis of the ATAC seq results (described above), indicating that the binding motifs that differed between these cells were associated with and may regulate those genes that similarly changed in expression. Transcription factors that bind to these motifs were also found to change expression in accordance with changes in motif accessibility (Figure 3F,G). Interestingly, those putative cis-regulatory elements that were accessible in both MG and progenitors, but that lost expression in mature MG (LOE/SA) are associated primarily with genes for cell cycle. These regions were enriched for the binding motif for the cell-cycle regulator E2F transcription factor (Figure 3D,E). This suggests that the differences between progenitor cells and MG in their cell proliferation are unlikely to be regulated by changes in chromatin accessibility.

### Ascl1 is Sufficient to Induce Neurogenesis in Immature MG

Our previous results showed that histone modifications may play a role in limiting the competence of MG to regenerate neurons: HDAC inhibition is necessary for Ascl1 to reprogram adult MG to neurogenic progenitors (Jorstad et al. 2017). Given the increased chromatin accessibility at progenitor genes of immature MG that we found at P8, we predicted that it might be possible to reprogram glia to a neuronal fate at P8. To test the hypothesis, we overexpressed Ascl1 in retinal progenitors and MG at various times during postnatal development. In order to trace the lineages of retinal progenitors and MG, we used a tamoxifen-inducible creER mouse driven by one of two promoters to activate a fluorescent reporter. For control mice, we used Glast-CreER:flox-stop-CC-GFP; mice received an intraperitoneal injection of tamoxifen to initiate the recombination at P0, P4, P8, or P12. The promoters are glial-specific in adult mice, but are expressed in retinal progenitors at P0. When cells were lineage traced at P0, the late-born types of neurons are present in the progeny: 72.6% of GFP+ cells had a photoreceptor fate, 22.7% were bipolar cells, and the remaining cells were MG (Figure 4A,C). These are similar to the ratios found when these cells that are birth-dated at this age in mice (Young 1985). By P4, the percentage of GFP+ neurons was significantly reduced with only 2.72% of cells having a bipolar cell fate, while the remainder throughout the retina were MG. At the far retinal periphery however, some retinal progenitor cells were still present at P4 and so some rods and bipolar cells were observed in this area (Figure S4). In mice older than P4, 100% of GFP+ cells were MG. From P8 and on, 100% of GFP+ cells were MG. These results are consistent with previous reports that neurogenesis in the retina ends after P4, with the exception of some neurogenesis at the periphery.

**Figure 4.**
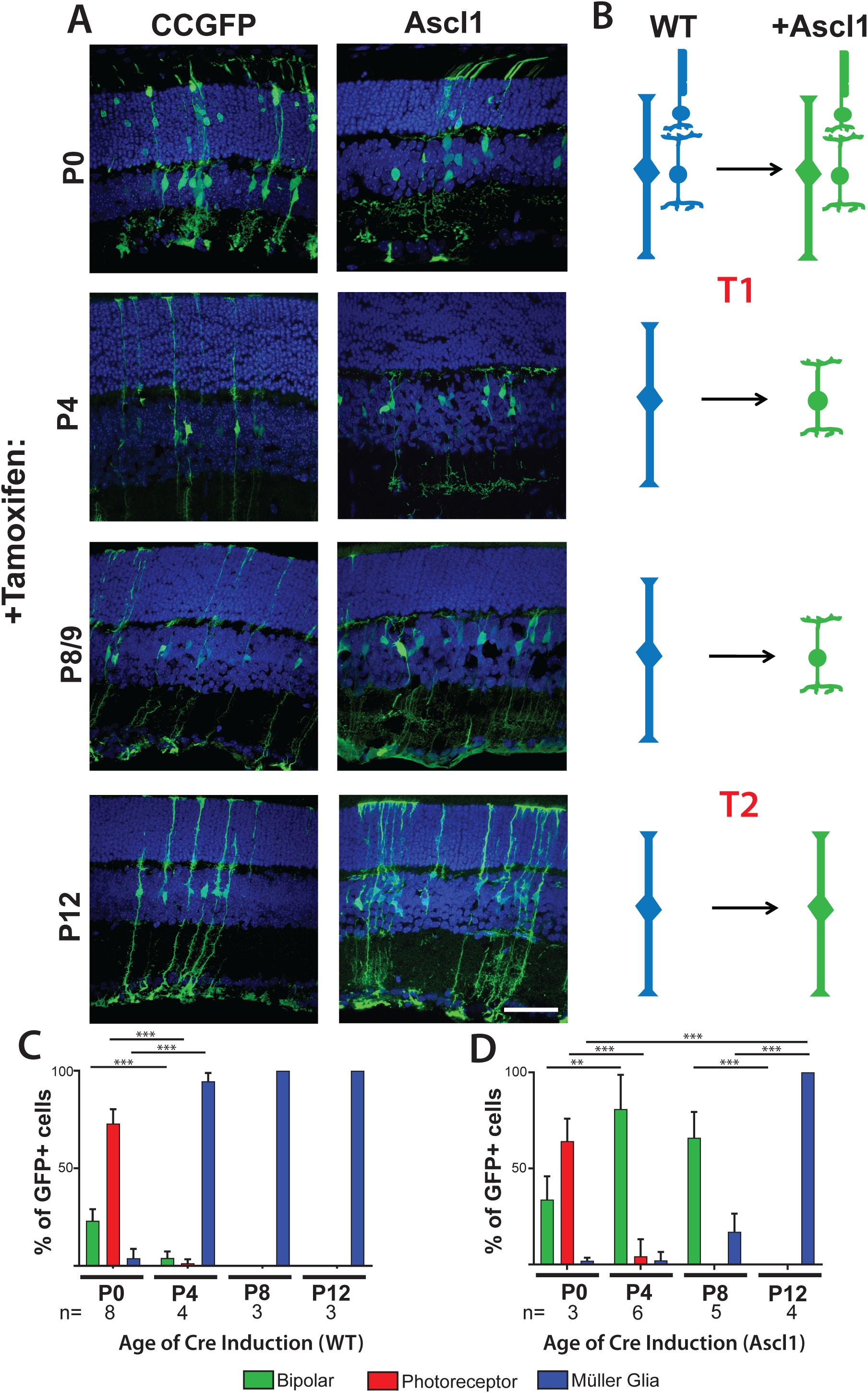
Ascl1 overexpression at young ages is sufficient to generate bipolar neurons. A. Cell tracing of progenitors and MG was induced with CC-GFP or Ascl1-ires-GFP on a Glast promoter at P0, P4, P8/9, and P12. All analyzed at P21. Scale bar 50 um B. Graphical summary of reprogramming observed with Ascl1. C. Cell counting quantification of GFP cell tracing in (A). D. Cell counting quantification of Ascl1 induction tracing in (A).

To determine whether immature MG can be reprogrammed to a neurogenic state with Ascl1 alone, we drove expression of Ascl1 at the same ages that we had used to trace WT cells. We used a previously described system for driving Ascl1 in MG: Glast-CreER;Flox-stop-LNL-tTA;TetOCMV-Ascl1-ires-GFP (Ueki et al. 2015; Jorstad et al. 2017). With the induction of Ascl1 expression at P0, ratios of GFP+ neurons were not significantly different from WT lineages. This suggests that the level of Ascl1 is expressed at sufficient levels in retinal progenitors to sustain neurogenesis, and that additional Ascl1 has little effect. At P4 and P8, however, over-expression of Ascl1 in immature MG has a dramatic effect on the cells: 75% and 65% of GFP+ cells, respectively, are now bipolar neurons, with few—if any—photoreceptor neurons. When we delay induction of Ascl1 to P12, however, 100% of GFP+ cells are now MG, and at this age, expression of Ascl1 alone is no longer sufficient to reprogram the MG to neurons, similar to results previously reported(Ueki et al. 2015; Figure 4A,D). These results show there is a rapid change in the competence of cells to generate rods with Ascl1-overexpression summarized in Figure 4B. Some major change (Transition [T]1) occurs at P4 in the progenitor cells as they become MG to restrict the ability of Ascl1 to generate rods. A second major change (Transition [T]2) appears to control the ability of MG to generate bipolar cells, and this occurs after P8. After this time, damage to the retina and inhibition of histone deacetylases is needed to induce neurogenesis from the MG.

### Retinal Neuron Chromatin Overlaps Similarly with Progenitor and MG

The changes in neurogenic competence that occur as cells transition from progenitor cells to MG even with forced Ascl1 expression might be due to changes in their epigenome. To determine if DNA accessibility might underlie these differences in (1) neurogenic competence, and (2) fate restriction to predominantly bipolar cell neurogenesis, we compared ATAC-seq data from progenitor cells, MG, rod photoreceptors (Hughes et al. 2017), and bipolar cells (Jorstad et al. 2017) (Figure 5A). We found that both progenitor cells and adult MG accessible regions overlapped with rods by approximately 19k peaks or 47.5% of glial/progenitor-accessible domains (Figure 5B,D). A similar analysis of bipolar cells and retinal progenitors or MG gave very similar results: bipolar cell accessibility overlapped with progenitor accessible regions by approximately 20k peaks (50% of progenitor domains), and with MG accessible regions by approximately 17k peaks (42.7% of glial domains) (Figure S6). For all overlaps, GO categories were similar between comparisons to progenitors or MG (Table S8). As Otx2 is involved downstream of Ascl1 in the generation of both cell types (Omori et al. 2011), we looked for central enrichment of this motif in both neuronal-specific and MG-shared accessible regions. For rods, the rod-specific accessible regions (ie. not present in the MG or the progenitor cells) are strongly centrally-enriched for Otx2 binding sites (Figure 5C,E, S6). We did not see a similar central enrichment for Otx2 in the accessible regions shared between rods and progenitor cells or rods and MG. A similar analysis of the overlapping and uniquely accessible regions for bipolar cells and either MG or retinal progenitors gave strikingly similar results: the accessible regions that are unique to bipolar cells and not shared with progenitor cells or MG show strong central enrichment for the Otx2 binding motif; however, regions shared among these cell types did not have this signature. Thus, the ability of Ascl1 to promote the bipolar fate is not due to MG having an epigenome that is more similar to bipolar cells than rod photoreceptor cells.

**Figure 5.**
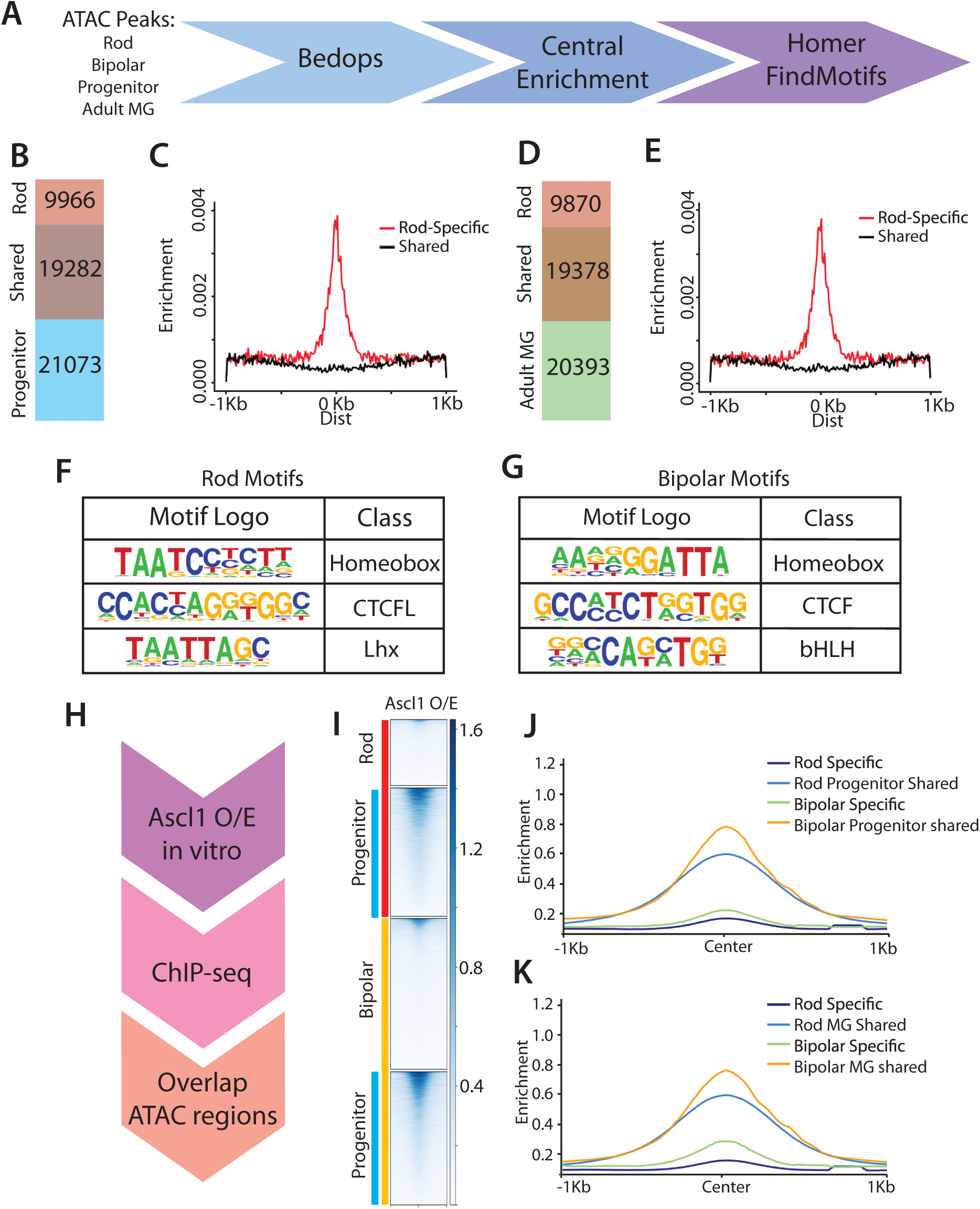
Ascl1 differentiates Bipolar fates from Rods. A. Analytical design comparing the accessible regions in neurons((Hughes et al. 2017; Jorstad et al. 2017) with ATAC-seq from progenitors and MG. B, D. Rod chromatin accessibility overlaps with both progenitor(B) and adult MG(D) accessibility by 19k regions. C,E. Otx2 predicted central enrichment is high in Rod-specific accessible regions, and low in regions shared with progenitors (C) and adult MG (E). F,G. Predicted motifs for rod-specific (F) and Bipolar-specific (G) accessible regions. H. Experimental design for Ascl1 ChIP-seq. I. Ascl1 ChIP-seq read density in Rod, Bipolar, and Progenitor accessible regions. J,K. Central enrichment for Ascl1 binding in rod, bipolar, and progenitor(J) or adult MG(K) accessible regions.

Overall motif enrichment in rod and bipolar neurons was additionally quite similar. Rod-specific regions—when comparing to either MG or progenitors—were enriched for homeobox, and CTCF domains, though in comparison to progenitors, rod photoreceptor cells may have an additional enrichment for bHLH binding motifs (Figure 5F, Table S2). Shared accessible regions in both comparisons showed enrichment for CTCF, Klf, and ETS binding motifs. Bipolar cell accessible regions that were shared with both progenitor cells and MG also showed enrichment for these TF binding motifs. Interestingly, in both comparisons, bipolar cell-specific accessibility demonstrated enrichment not only for homeobox domains, but also for proneural bHLH domains (Figure 5G). The presence of bHLH accessibility in bipolar cells may underlie the fact that MG primarily generate bipolar cells with the overexpression of Ascl1.

### Ascl1 Binding Sites are Enriched in Bipolar Regions

These results imply that bHLH transcription factors such as Ascl1 may play a specific role in directing MG to a bipolar cell fate. We next looked at predicted central enrichment of Ascl1 in these neuronal populations. Rod-specific accessible regions demonstrated some central enrichment of predicted Ascl1 binding motifs, though the overall enrichment did not appear to be notably elevated from regions with accessibility shared with progenitors or MG. By contrast, bipolar cell accessible regions had distinct central enrichment of predicted Ascl1 binding sites, which is also found in shared accessible regions, though to a lesser extent (Figure S6).

Because the role of Ascl1 appears to be a key difference between bipolar cell and rod photoreceptor cells, we took a closer look at its binding in accessible and inaccessible domains. Casey et al 2018 recently demonstrated that ES cell reprogramming with Ascl1 revealed a particular class of accessible regions with repeated Ebox motifs (Casey et al. 2018). Similar to that study, we find that those regions of the DNA that are inaccessible in MG or progenitors, but bind Ascl1 when over-expressed (pioneered sites) had on average 4.7 E-boxes per peak, while other Ascl1-bound sites in MG have fewer than 2 E-boxes on average (Figure S7). We additionally assayed E-box number in Rod and Bipolar accessible domains, but found no difference in average number of E-boxes per peak (DNS).

In addition to this predictive analysis, we compared accessibility domains using Ascl1 ChIP-seq from both P0 retinas as well as from Ascl1 overexpression in cultured MG (Figure 5H). While developmental Ascl1 largely binds in shared and progenitor accessibility domains as well as some non-glial sites, ectopic Ascl1 has a strong distinct niche in Adult MG and in so-called “radical” binding sites (Figure S7). In observation of regions predicted to be enriched in Ascl1 binding in retinal neurons, we found that Ascl1 also binds more strongly to bipolar accessible domains as compared to rod accessible domains (Figure 5I). Tag density was highest for progenitor and glial-shared accessible regions, but there was distinct central enrichment for Ascl1 binding in bipolar-specific accessible domains (Figure 5J,K). Binding of Ascl1 ectopically and Ebox patterning in novel accessible domains demonstrate that expression of this TF in MG may further direct fate decisions more towards bipolar interneurons than photoreceptor neurons.

## Discussion

In this study, we have explored the epigenetic profile of retinal progenitors and MG alongside their gene expression profile as these cells change in their neurogenic potential. We have demonstrated epigenetic evidence of putative cis-regulatory elements that change in accessibility through development and potentially regulate the maturation and neurogenic potential of the MG. Furthermore, we have shown that the intermediate profile of immature MG allows for improved reprogramming to bipolar neurons, thus revealing transition states in the neurogenic potential of immature mammalian glia.

In order to explore the potential mechanisms for regulation of epigenetic regenerative capacity in the mammalian MG, we performed ATAC-seq on postnatal neurogenic retinal progenitors as well as on developing and mature MG. We found that the epigenomic landscape of progenitors and mature MG were very similar, with approximately 60% of progenitor accessible domains shared with mature MG, consistent with similarities in expression between progenitors and MG (Jadhav et al. 2009; Nelson et al. 2011). These shared regions include glial-expressed genes as well as progenitor-specific genes and were particularly enriched around promoter regions. Cell-type specific accessible regions were enriched in intronic and intergenic regions of the genome, consistent with previous evidence that enhancers may be more dynamic during development(Aldiri et al. 2017).

Accessible regions that were enriched in the retinal progenitors were specifically associated with developmental and neurogenic genes. This was reinforced by expression analysis, demonstrating that progenitor-specific genes, enriched in GO categories associated with early development and neurogenesis, showed more than 2 logFC compared with MG. Many regions of accessibility that were reduced or lost as the MG mature were associated with these same genes that lost expression during postnatal development, reinforcing that accessibility correlates with expression. Motif enrichment analysis of the progenitor-specific accessible domains revealed specific enrichment in bHLH binding domains. This class of transcription factor is well-characterized as part of the retinal developmental process, though there are many variations of E-box motifs in the genome that are relatively specific to different bHLH domains. Aydin et. al 2019 recently described variations of E-box specificity that contribute to neuronal subtype (CAGSTG for Ascl1 and CAKATG for Neurog2) (Aydin et al. 2019); we found a nonspecific Ebox enriched by HOMER, consistent with progenitor cell potential to generate multiple types of neurons (both excitatory and inhibitory), with the fourth position in the motif being equally probable to be A or C. However, specific assaying for specific Ebox motifs demonstrates a preference for the Ascl1 motif throughout. The loss seen in bHLH binding motifs in the transition from progenitors to MG confirms that these transcription factors reflects the decline in expression of Ascl1 and related bHLH TFs as progenitors transition to MG.

As MG mature, the cells increased in both accessibility and expression associated with genes important for glial function. There was a correlation between increases in nearby accessible chromatin and genes that increased in expression during glial maturation. Young MG in particular had a unique intermediate profile. Glial genes that were highly increased in expression in adult MG gain accessibility before P8, but this is not accompanied by a similar drop in progenitor gene expression.

Cis-regulatory accessibility continued to change and develop during MG maturation. While many enriched motif classes were common between decreasing and increasing accessibility, the NFI binding motif was uniquely enriched in the developing glia. NFI domains were similarly enriched surrounding genes that increase expression and gain accessibility, indicating a putative role in the development and maturation of the glial fate. These NFI domains were enriched in accessible regions even in the younger P8 MG, consistent with evidence for an intermediate profile of immature MG. NFI transcription factors are relevant to the developing glia as they are well known for their role in CNS glial development (Shu et al. 2003; Heng et al. 2015; Deneen et al. 2006). In addition, NFIa/b/x are expressed in retinal progenitors and MG, and conditional knockouts of these transcription factors in the developing retina are associated with defects in gliogenesis and the production of bipolar neurons (Clark et al. 2019). As progenitors develop, there is an acquired association of NFI motifs with Lhx2 binding sites, which may be related to guiding the neurogenic potency of late retinal progenitors (Zibetti et al. 2019).

Though RNA-seq and ATAC data demonstrate epigenomic changes that co-occured with changes in gene expression, not all gene expression changes were associated with changes in accessibility. This is best demonstrated with cell cycle and proliferation-related genes. These regions are associated with E2F binding motifs, consistent with their roles in regulating mitotic proliferation (Wu et al. 2001; Stevaux and Dyson 2002). Expression of genes associated with the mitotic cell cycle declined as progenitors withdrew from the cell cycle. However, regions of accessibility associated with these genes changed little in their accessibility between progenitors and MG. This suggests that the loss of proliferative capacity in maturing MG may not be limited by epigenetic accessibility.

Our results showed that the immature (P8) MG have an epigenome intermediate between the progenitors and adult MG. Since efficient reprogramming of mature MG requires a combination of Ascl1 and HDAC inhibition, we postulated that the P8 MG might be more efficiently reprogrammed to a neuronal fate with Ascl1 alone. We found that induction of Ascl1 alone in immature MG was indeed capable of inducing a neuronal/bipolar cell fate, consistent with our hypothesis that epigenomic changes in accessibility limit mature MG from regeneration. Interestingly, the P8 MG already appeared to be fate restricted with respect to the types of neurons produced from Ascl1 over-expression, with two distinct transition periods. In newborn mice, lineage tracing and birthdating studies have shown that progenitors generate three types of neurons: rods, bipolar cells and amacrine cells (Young 1985); however, early in MG development (eg. P4), the MG cells lose their ability to generate photoreceptors and amacrine cells (T1), even with Ascl1 over-expression. Somewhat later in MG development (eg. P10) as the cells mature, they lose their ability to generate bipolar cells from Ascl1 over-expression alone, though the addition of HDAC inhibitors and injury can restore their neurogenic potential (Ueki et al. 2015; Jorstad et al. 2017).

Do changes in accessible chromatin account for the bipolar fate restriction that occurs during the T1 transition? To address this question, we compared the open chromatin landscape of bipolar cells and rod photoreceptors to determine the degree of similarity between these neuronal cell types and MG or progenitors. Overall, the shared accessible regions between progenitors and either type of neuron (rod or bipolar cell) were very similar to the shared regions between MG and these types of neurons. Moreover, the newly accessible sites in rods or bipolar cells were not present in either progenitors or MG. Thus, the degree of similarity in accessible chromatin between progenitors or MG, on the one hand, and rods or bipolar cells, on the other, are not sufficient to explain the bias in bipolar cell generation from MG. Nevertheless, in comparing neuron-specific accessible regions from rods and bipolar cells, we found that bipolar cell-specific accessible regions were more highly enriched for bHLH motifs than rod-specific accessible regions. Also, over-expression of Ascl1 in MG results in Ascl1 binding to bipolar cell-specific accessible regions over rod-specific regions. Similar enrichment for Ebox accessibility in a variety of bipolar cells was also shown by Murphy et al (Murphy et al. 2019), and this may help to explain why the overexpression of Ascl1 in MG preferentially generates bipolar cells *in vivo*, though the factor is necessary for the generation of both bipolar cells and photoreceptors during development.

In sum, we have characterized the ways in which the accessible chromatin landscape changes as retinal progenitors differentiate into MG. Developing MG lose expression of neurogenic genes and accessibility of related cis-regulatory elements, and gain accessibility of glial-defining NFI binding sites. However, young MG demonstrate intermediate profiles. By P8, MG demonstrate early gains in NFI binding sites, while retaining progenitor-like expression and accessibility. This intermediate profile translates into neurogenic potential: prior to P12 overexpression of Ascl1 alone is sufficient to induce neurogenesis in MG. The affinity for bipolar neurons appears to be in part due to the preference of Ascl1 for binding to bipolar cell specific accessible regions, and not due to inherent overlaps in MG chromatin accessibility with bipolar cells vs rods. Overall, our results show unique transition states in the development of glial cells that restrict their neurogenic potential, which correlate with changes in the epigenome.

## Methods

### Mice

All mice were housed at the University of Washington and treated with protocols approved by the University of Washington’s Institutional Animal Care and Use Committee. Mice expressed Cre-recombinase under the *Glast* promoter from Jackson Labs with *Rosa-flox-stop-tTA* (Jackson labs) in combination with either *tetO-mAscl1-ires-GFP* (M. Nakafuku University of Cincinnati) or CCGFP. Mice having EGFP knocked-into the Sox2 open reading frame were obtained from Jackson Laboratories (Stock: 017592) and bred to generate P2 litters(Arnold et al. 2011). *Rlbp1CreERT2* mice were crossed to *R26*-*stop*-*flox*-*CAG-tdTomato* mice (Jackson Labs, also known as Ai14; 129SvJ background). Mice of both sexes were used for this study and analyzed together in their respective treatment groups.

### Injections

Intraperitoneal injections of 50 μl of tamoxifen (Sigma) at 100 mg/kg in corn oil were given to induce expression of Ascl1 and GFP. Tamoxifen was administered once for P0 or P4 induction, for 2 consecutive days in mice aged P6-P9, or 4 consecutive days for older ages.

### IHC and cell counts

For lineage tracing with GFP and Ascl1 induction, animals were euthanized and the eyes removed for dissection and removal of the cornea and lens. Eyes were then fixed in 4% PFA in PBS for 1 h before being transferred to a 30% sucrose in PBS solution and kept overnight at 4 °C. Eyes were then frozen at −80 °C in O.C.T. (Sakura Finetek) and sectioned to 14μm by cryostat (Leica). Slides were incubated at room temperature in blocking solution (10% normal horse serum, 0.5% Triton X-100, in PBS) for 1 h. Slides were then incubated overnight at 4 °C in primary antibody diluted in blocking solution. Slides were then washed three times in PBS for 15 minutes each before a 1 h room temperature incubation in secondary antibodies (Life Technologies) in PBS. Slides were washed once before being incubated 5-10 minutes with 1:10,000 DAPI (Sigma) in the dark. At this point, slides were washed three times in PBS and coverslipped with Fluoromount-G (SouthernBiotech). Primary antibodies: goat anti-Sox2 (Sanda Cruz, 1:500), mouse anti-HuC/D (Invitrogen, 1:100), chicken anti-GFP (Abcam, 1:500), goat anti-Otx2 (R&D Systems, 1:100), rabbit anti-Recoverin (Millipore, 1:1000).

Section imaging was performed using an Olympus Fluoview confocal microscope, and random fields throughout the retina were sampled for cell counts. Cell types were identified and counted by localization within the retina, cell morphology, and marker co-staining. Significance values between treatments were determined by one-way ANOVA with a post-hoc tukey test or by t-test.

### FACS

#### FACS protocol according to (Wohl and Reh 2016)

Retinas were isolated via dissection away from surrounding tissues and then washed in PBS. Fluorescence was confirmed via live imaging under an inverted fluorescent microscope (Zeiss). Retinas were then incubated on a nutator in Papain and DNase I for 10 min at 37 °C. Retinas were then triturated to generate a single-cell suspension and transferred to a tube containing an equal volume of Ovomucoid. The suspension was then spun down at 300*g* for 10 min at 4 °C. Cells were resuspended in a solution of 100:1:1 Neurobasal:DNase:Ovomucoid, and passed through a 35 μm filter before sorting using a BD FACSAria III Cell Sorter (BD Biosciences). Gating was performed to isolate intact cells from debris and to isolate positive fluorescent glial or progenitor cells. Progenitor cells were isolated from Sox2+ Amacrine cells by removing the higher fluorescent neuronal population. Positive fractions containing fluorescently labelled MG were then spun down at 300*g* and resuspended for the appropriate assay.

### ATAC

Purified cells from live-cell FACS were input into a 15 μl transposase reaction with an input of 100k cells in a protocol modified from the Greenleaf lab (Buenrostro et al. 2015). Transposition was carried out with reagents from the Nextera DNA Sample prep kit: 7.5 μl 2X TD Buffer, 0.75 μl Tn5 Transposases, and nuclease-free water to 15 μl. The reaction was mixed and incubated at 37 °C for one hour before being purified with the Qiagen Reaction Cleanup Minelute kit and eluted into 10 μl. Libraries were prepared through subsequent PCR using Illumina Nextera kit (Cat. No. FC-121-1030) using a test qPCR output to estimate the number of cycles necessary to properly amplify the library. Amplified libraries were purified with the Qiagen PCR cleanup minelute kit and eluted into 20 μl. Library QC was performed using gel electrophoresis, and quantitated on a Qubit 3.0 Fluorometer with the dsDNA HS Assay kit and A260/280 and A260/230 checked by nanodrop before sending for Illumina NextGen sequencing on a Next Seq 500 in rapid mode employing a paired-end 50 base read length sequencing strategy (Seattle Genomics). Adapter and barcode sequence were removed from the reads and low-quality sequences (Phred score <33) were removed using Trim Galore (Krueger 2015). Remaining reads were mapped using Bowtie2 (Langmead and Salzberg 2012), marking duplicate reads with Picard (http://broadinstitute.github.io/picard/), and removing reads using Samtools (Li et al. 2009). Alignment data was normalized for coverage using a custom R script (https://rpubs.com/achitsaz/98857) and visualized using the Integrated Genomics Viewer(Robinson et al. 2011)

### Peak calling and comparison

Peaks were called using HOMER (Heinz et al. 2010) findPeaks dnase style with a minimum distance of 415 and size of 150. Bedops -e 1 and -n 1 functions were used to compare peak files for binary peak differences (Neph et al. 2012). For differential accessibility comparisons, the R package EdgeR (Robinson et al. 2010) was used to compare all peak regions between two ATAC samples against the reads of each file as previously described. The RSubread (Liao et al. 2019)function featureCounts was used to generate a matrix of counts per million across all peaks. The counts matrix was filtered against low-reads rows and in the case of any sample that is more deeply sequenced than another, the EdgeR (Robinson et al. 2010) function thincounts was used to thin one sample randomly to the level of the lower depth of sequencing. Dispersion, fitting and differential signal testing were performed using negative binomial generalized linear models as specified in the edgeR guide. Cumulative differences in accessibility at each gene were calculated as the sum of the fold differences of all peaks nearest to each gene. Peaks of interest were identified by selecting for those with a log2FC above 2, and the top 1k peaks up and down were selected by log2CPM. The peakIDs for each of these regions were used to generate new peak files and perform further analysis.

### Ascl1 Chromatin Immunoprecipitation-Sequencing (ChIP-Seq)

P0 retinas or cultured, post-natal day 12, Müller glia (+/- Ascl1 overexpression, rtTA germline:tetO-Ascl1-ires-GFP mice ± doxycycline) were digested with papain/DNase to single cells and fixed with 0.75 % formaldehyde for 10 minutes at room temperature. Sonication was performed with a probe sonicator (Fisher Scientific): 12 pulses, 100 J/pulse, Amplitude: 45, 45 seconds cooling at 4 °C between pulses. Immunoprecipitation performed with 40 μL anti-mouse IgG magnetic beads (Invitrogen Cat: 110.31) and 4 μg mouse anti-MASH1 antibody (BD Pharmingen Cat: 556604) or 4 μg mouse IgG against chromatin from 5 million cells per condition according to Diagenode LowCell Number Kit using IP and Wash buffers as described in (Castro et al. 2011). Libraries were prepared with standard Illumina adaptors and sequenced to an approximate depth of 36 million reads each. Sequence reads (36 bp) were mapped to the mouse mm9 genome using bwa (v 0.7.12-r1039). Merging and sorting of sequencing reads from different lanes was performed with SAMtools (v1.2). The HOMER software suite was used to determine and score peak calls (‘findPeaks’ function, v4.7) as well as motif enrichment (‘findMotifs’ function, v4.7, using repeat mask). For STATi and control Ascl1 ChIP-seq, reads were aligned to the mm10 genome using Bowtie2. The .sam files were converted to sorted .bam files using SAMtools. MACS2 was used to call peaks with default settings using the broad peaks annotation. Peak overlap analyses were performed using Bedops. The control Ascl1 ChIP-seq .bam file was downsampled by a factor of 0.69 to normalize the number of mapped reads over the common peaks found between treatment and control samples. This downsampled .bam file was used for all analyses. Differential accessibility analysis in Ascl1 ChIP-seq peaks was determined using edgeR as detailed in the edgeR user guide.

### Bulk RNA seq

For RNA-Seq, FAC-sorted cells were resuspended into Qiazol and RNA was extracted using the Qiagen RNeasy MinElute kit. Samples were tested for QC using nanodrop, and sent for sequencing. 500 ng per sample (50 ng μl^−1^) was sequenced on an Illumina HiSeq and reads that passed Illumina’s base call quality filter were mapped to mm10 using TopHat v2.0.12. To generate counts for each gene using htSeq-count v0.6.1p1, in “intersection-strict” overlap mode, genes with zero counts across all samples were removed, and data normalized using edgeR v3.12.0. Further analysis was done using Bioconductor and R (version 3.2.3).

EdgeR was used to compare reads across RNA-seq samples according to the published manual guide for exactTest(Robinson et al. 2010). Prior to testing, samples with higher reads were thinned to the level of other samples using thinCounts. Dispersion, fitting and differential signal testing were performed using negative binomial generalized linear models as specified in the edgeR guide. Differential expression was filtered to generate the top 1k genes up and down across development by selecting for genes with a logFC greater than two, and top genes of this category were selected by highest log2CPM. The top genes were clustered using hclust to perform ward D2 agglomeration with Euclidian distances. Genes were filtered against annotated ATAC peaks by dplyr.

### Cell culture and expression

#### From (Pollak et al. 2013)

MG from postnatal day (P)12 mice were cultured [Neurobasal + N2, epidermal growth factor (EGF), 10% fetal bovine serum (FBS)] as previously described with 1 μM 5-ethynyl-2′-deoxyuridine (EdU). Lentiviral particles were added in Optimem (Gibco) or neural medium (Neurobasal + N2, B27, 1% tetracycline (tet)-free FBS), and 3-6 hours later medium was replaced. hBDNF (R&D Systems, 10 ng/ml), bFGF (R&D Systems, 100 ng/ml) and rGDNF (R&D Systems, 10 ng/ml) were added for longer cultures. 4-Hydroxytamoxifen (4-OHT; Sigma) was included where indicated at 10 μM.

### Homer annotatePeaks.pl

For basic annotation of ATAC peaks, the Homer function annotatePeaks.pl was used (Heinz et al. 2010). While in most cases, peak annotation to genes was accomplished with the online tool GREAT (McLean et al. 2010), annotating each region for closest gene, the basic usage was used: annotatePeaks.pl <peak file> mm9 > output.txt

For stats on genome location:

annotatePeaks.pl <peak file> mm9 -annStats > output_stats.txt

For annotation of specific motifs, a position weight matrix was either taken from the homer database, from the findmotifs function, or was generated using seq2profile.pl in Homer, and was used in annotatePeaks as such:

annotatePeaks.pl <peak file> mm9 -m pwm.motif > output_motif.txt

However, for the generation of lineplots, a table of 10bp bins for motif enrichment were generated for graphing with ggplot2:

annotatePeaks.pl <peak file> mm9 -m pwm.motif -size 2000 -hist 10 > output_plot.txt

For motif co-occurrence:

<u>annotatepeaks.pl</u> <peak file> mm9 -size 2000 -hist 20 -m <motifs of interest> -cpu 10 -matrix

fileout > fileout.motif.freq

### Homer Motif discovery

To discover enrichment of predicted DNA binding motifs for further analysis, we employed the Homer function findMotifsGenome.pl using suggested basic usage settings. From there, we identified top motifs from the homerResults output

### Gene Ontology

For annotation of gene ontology (GO) categories, we used the Bioconductor package GOstats (Falcon and Gentleman 2007). Gene lists from ATAC and RNAseq were input to a hyperGTest with a p value cutoff of 0.001 for Biological Process ontology categories. The top 20 GO terms were plotted in ggplot2. To acquire genes of interest from specific GO categories, we found annotations of the category ID on the Jax Mouse Genome Informatics database and subset our original genelist based on the genes in each GO category.

### Heat maps and plots (deeptools)

To generate heat maps and lineplots of ATAC and ChIP read enrichment, we used deeptools (Ramírez et al. 2016) computeMatrix reference-point to calculate enrichment scores by region along a bed file:

computeMatrix reference-point -S <bigwig files> -R <bed files> --referencePoint center -a 1000 -b 1000 –skipZeros – MAT_file.tab.gz

This is then plotted with the command:

plotheatmap -m MAT_file.tab.gz -out HM_file.png –colorMap Blues –missingdatacolor 1.0

Or, to plot the lineplots only:

plotProfile -m MAT_file.tab.gz -out Plot_file.png –yMax 3.0

## Acknowledgements

The authors acknowledge the following funding sources for supporting this work. Grant #TA-RM-0614-0650-UWA from the Foundation Fighting Blindness to T.A.R., NIH NEI 1R01EY021482 to T.A.R., Allen Distinguished Investigator Award (Paul G. Allen Family Foundation) to T.A.R., an NSF Fellowship to M.S.W. (DGE-0718124), Scholarship Wo 2010/1-1 from Deutsche Forschungsgemeinschaft (DFG) and the SUNY Empire Innovation Program Grant to S.G.W. and NIH training grant T32HD007183 and PHS NRSA T32GM007270 from NIGMS to L.S.V.

We thank members of the Reh and Bermingham-McDonogh laboratories for their review and valuable discussion regarding the manuscript. In particular, we thank Olivia Bermingham-McDonogh, Levi Todd, Marcus Hooper, and Brent Wilkerson for their comments on the manuscript. Lastly, the authors thank M. Nakafuku (Cincinnati Children’s) for the tetO-Ascl1-ires-GFP mice.

## Author Contributions

LSV and TAR conceived of all experiments and analyses. LSV conceived of and performed all ATAC experiments and analyses. SGW and LSV performed FACS and RNA-seq experiments, and LSV performed all analyses. MSW conceived of and performed culturing and Ascl1 ChIP-seq experiments. KC and LC performed cell tracing and in vivo Ascl1 experiments and cell counts. LSV and TAR wrote the manuscript with input from co-authors.

